# Lessons learned while evaluating how pairing aversive experiences with same or different sex social stimulus mice affects subsequent social engagement

**DOI:** 10.1101/2024.08.12.607663

**Authors:** Jasmin N. Beaver, Marissa M. Nicodemus, Isabella R. Spalding, Lauren R. Scrimshaw, Sohini Dutta, Aaron M. Jasnow, Lee M. Gilman

## Abstract

Mice offer a wealth of opportunities for investigating brain circuits regulating multiple behaviors, largely due to their genetic tractability. Social behaviors are translationally relevant, considering both mice and humans are highly social mammals, and human social behavior disruptions are key symptoms of myriad neuropsychiatric disorders. Stresses related to social experiences are particularly influential in the severity and maintenance of neuropsychiatric disorders like anxiety disorders, and trauma and stressor-related disorders. Yet, induction and study of social stress in mice has disproportionately focused on males, influenced heavily by their inherent territorial nature. Social target-instigated stress (i.e., defeat), while ethologically relevant, is quite variable and predominantly specific to males, making rigorous and sex-inclusive studies challenging. In pursuit of a controllable, consistent, high throughput, and sex-inclusive method for social stress elicitation, we modified a paradigm to train male and female F1 129S1/SvlmJ × C57BL/6J mice to associate (via classical conditioning) same or different sex C57BL/6J targets with a mild, aversive stimulus. While further paradigm optimization is required, social interaction testing 24 h after conditioning indicates males socially conditioned better to male targets by exhibiting reduced social interaction, whereas females appeared not to form social stimulus associations. Serum corticosterone levels inversely corresponded to social avoidance after different sex, but not same sex, conditioning, suggesting corticosterone-mediated arousal influences cross-sex interactions. These rigorously controlled null outcomes align with past pursuits’ limited success in creating a sex-inclusive social stress paradigm.

**Significance Statement:** Validated paradigms to study social stress in female mice, and across sexes, are needed. We modified a published male mouse protocol by using classical conditioning to pair an aversive stressor with a target. Our goal was to create a uniform, cross-sex, high-throughput social stress technique to advance future research. Though our modified paradigm requires future improvements, we did acquire evidence that males can be socially conditioned in this way, and female same sex social engagement can be attenuated by a preceding non-social aversive experience. These null findings, while not achieving our goal, provide useful information to advance future sex-inclusive social stress investigations.

## Introduction

Social interaction behavior is a cross-mammalian phenomenon seen in humans, non-human primates, rats, hamsters, and mice, among other animals (Bloomberg et al., 1994; Jasnow and Huhman, 2001; Kamps et al., 2001; Leung, 2011; Monfils and Agee, 2019; Lee et al., 2020). Social interaction behavior shifts can be informative of the physical and/or emotional state of the mammal (Ago et al., 2014; Wilson and Koenig, 2014; Venniro et al., 2018). In humans, typical social interaction behavior disruptions are characteristic of numerous neuropsychiatric conditions including trauma and stress-related disorders, mood and anxiety disorders, and autism spectrum disorder. (American Psychiatric Association, 2022).

Rodent studies examining social behavior and socially associated stressors within various contexts have advanced identification of regulating neural pathways, and assisted development of new therapeutic approaches (Jasnow and Huhman, 2001; Coccurello et al., 2009; Smith and Wang, 2012; Gilman et al., 2015; Caldwell et al., 2017; Bergamini et al., 2018; Donovan et al., 2020; Sahani et al., 2022; Venniro et al., 2022). However, social behavior research in the genetically tractable mouse species (*Mus musculus*) has been relatively limited in studying behavioral and neurophysiological consequences of aversive social interactions in *female* mice (Kondrakiewicz et al., 2019; Kuske and Trainor, 2021). This is largely due to capitalization of male-specific territorial aggression in most rodent social conditioning paradigms (Blanchard et al., 2001; Gilman et al., 2015; Latsko et al., 2016; Peña et al., 2019; Lopez and Bagot, 2021; Furman et al., 2022; Lyons et al., 2023; Pantoja-Urbán et al., 2024). A breadth of social stress techniques have been employed in hamsters, mice, and rats (Miyashita et al., 2006; McCormick et al., 2007; Bernberg et al., 2008; Kercmar et al., 2011; McCormick et al., 2013; Iñiguez et al., 2018; Sterley et al., 2018; Lee et al., 2020; Furman et al., 2022; Pan et al., 2023). Of these, the most prevalent paradigm in mice is that of social defeat (both acute and chronic) (Jasnow and Huhman, 2001; Jasnow et al., 2015; Golden et al., 2011; Dulka et al., 2015; Gilman et al., 2015; Latsko et al., 2016; Bonnefil et al., 2019; Wang et al., 2021). Aside from the male-centric nature of social defeat, additional concerns of reproducibility arise from the inherently variable range of aversive social experiences each ‘defeated’ experimental mouse encounters. Such variability, mostly beyond experimenter control, plus the single sex bias of social defeat, left us wanting to explore an improved paradigm. For this, we paired controllable, uniform, aversive unconditioned stimuli with the presence of a social stimulus (conditioned stimulus; target) to study aversive social conditioning *across sexes* in mice using a higher throughput approach (requiring one conditioning day, rather than multiple days-weeks).

To accomplish this, we modified a paradigm previously utilized in male mice, involving manual administration of an aversive stimulus (mild foot shock) selectively when a male mouse actively investigated a target, with the goal of attenuating subsequent social engagement (Toth et al., 2012; Zoicas et al., 2014; de la Zerda et al., 2022; Grossmann et al., 2024). Rather than continuing this operant-style approach, where a mouse’s behavior dictates an outcome, we shifted to a classical conditioning approach in which shock delivery is standardized across individual mice. We anticipated experimental mice would associate the presence of the target with the aversive unconditioned stimulus. Additionally, all mice would receive the same number of shocks, instead of variable numbers based upon behavior.

Here, we evaluated how this paradigm affected social engagement and fear behaviors *across sexes* after mice were exposed to a novel target. Specifically, mice were socially conditioned with an aversive stimulus (mild foot shock; stress) when in the presence of their assigned target, independent of investigative behavior (Social Stimulus + Stress group). Then, mice were tested for social engagement and fear behavior in a separate environment with their assigned target present, followed by testing of freezing in the conditioning environment in the absence of any target. The former test enabled evaluation of both social and fear behaviors, while the latter test assessed contextual memory sans social stimuli. These behaviors were tested under circumstances when the experimental and target mice were the same sex, and when they were different sexes. We hypothesized Social Stimulus + Stress mice would exhibit reduced social engagement (indexed as time spent directly adjacent to the target enclosure) when tested with their assigned target in a novel environment, regardless of if the Social Stimulus + Stress and target mice were the same or different sexes. Our broad goal was to develop a paradigm that would facilitate rigorous future investigations of social behavior across mouse sexes, which are currently underrepresented in literature (Kuske and Trainor, 2021).

## Methods

### Mice

Hybrid F1 offspring (i.e., 129SB6F1/J) of both sexes resulting from pairing female 129S1/SvlmJ (RRID:IMSR_JAX:002448) and male C57BL/6J (RRID:IMSR_JAX:000664) mice (hereafter experimental mice), and male and female C57BL/6J mice (hereafter targets), all ≥9 weeks old or older, were group-housed (two to five per cage) within sex. Experimental mice were hybrid F1 offspring of 129S1/SvlmJ and C57BL/6J mice because these two strains are the most prevalent among transgenic mice, and because hybrids provide broader generalizability (Kumar et al., 2021; Keady et al., 2022). Such hybrids are used for a variety of physiology and neuroscience research purposes (e.g., (Curley et al., 2010; Kelley et al., 2011; Agrimson et al., 2017; Niibori et al., 2020; Correia et al., 2024)). Targets were C57BL/6J mice housed in a separate room from experimental mice. C57BL/6J mice were used as targets to ensure they were entirely novel to experimental mice through separate housing; because C57BL/6Js are often used as the primary or background strain for testing social behaviors in mice (see reviews (Laman-Maharg and Trainor, 2017; Varholick et al., 2021; Takahashi, 2025)); and because our F1 breeding was limited to generation of experimental mice. All mice had *ad libitum* access to food and water in rooms maintained on a 12:12 light/dark cycle with lights on at 07:00 local standard time (i.e., Zeitgeber time 0), and temperature maintained at 22 ± 2°C. All mice were fed LabDiet 5001 rodent laboratory chow (LabDiet, Brentwood, MO) and were kept on 7090 Teklad Sani-chip bedding (Envigo, East Millstone, NJ) in cages containing Nestlets (Ancare, Bellmore, NY) and huts (Bio-Serv, Flemington, NJ) for enrichment. Experiments were approved by Kent State University’s Institutional Animal Care and Use Committee, and adhered to the National Research Council’s Guide for the Care and Use of Laboratory Animals, 8^th^ Ed. (National Research Council, 2011).

### Social Conditioning Paradigm

#### Timeline

The entire paradigm spans four consecutive days. Procedures for each individual day are outlined below in order. On Days 0 and 1, experimental mice underwent different experimental manipulations according to their specific treatment group. On Days 2 and 3, all experimental mice experienced identical testing conditions regardless of their treatment group (Fig. 1).

**Figure 1.**
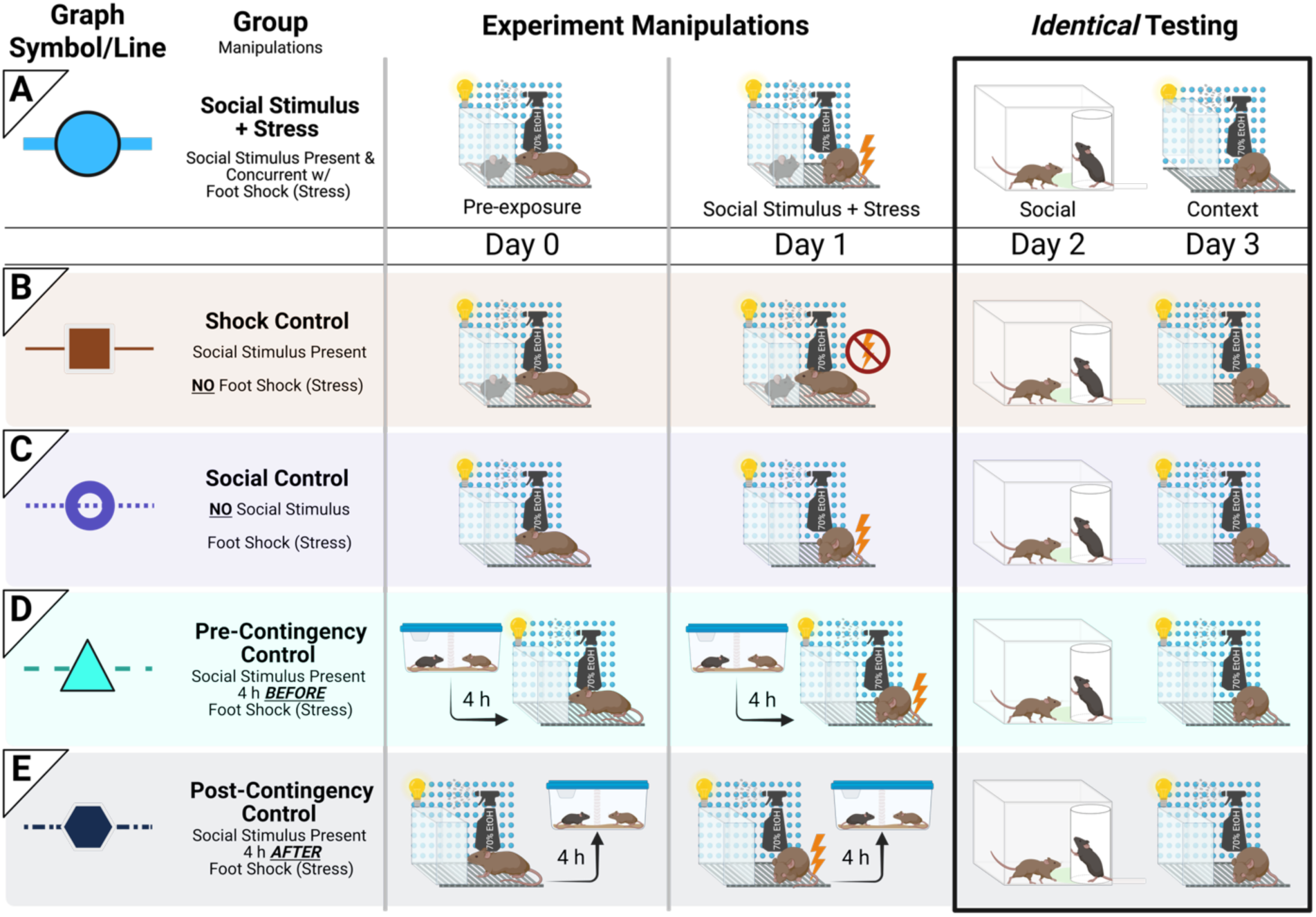
Study timeline and treatment groups. Five groups were used in this study for each experiment (same sex experiment and different sex experiment): one socially conditioned (Social Stimulus + Stress) group and four Control groups. Day 0 was for pre-exposure; Day 1 was when social conditioning occurred; Day 2 tested for social engagement; Day 3 tested for context-specific freezing behavior. On Day 0, Social Stimulus + Stress mice (A; above horizontal lines) encountered their assigned target in the social conditioning context, where olfactory, visual, and auditory cues could be exchanged while tactile interactions were minimized. No aversive stimulus was applied on Day 0, but on Day 1, experimental mice and their assigned target were returned to this context. Social Stimulus + Stress (but not target) mice then experienced five mild foot shocks to associate their target with that aversive experience. Social Stimulus + Stress mice were then tested for social engagement with that same target on Day 2, followed by testing for context fear behavior on Day 3 in the absence of their assigned target. Each Control group (below horizontal lines) was composed of different mice. Shock Control (B) mice underwent the same exact procedure as detailed for the Social Stimulus + Stress group, except no foot shock was administered on Day 1. Social Control (C) mice similarly experienced the same procedure as mice in the Social Stimulus + Stress group, save that Social Control mice did not encounter any target until Day 2 for testing of social engagement. Pre-Contingency Control (D) mice encountered their target four h before pre-exposure and social conditioning, meaning no target was present for Pre-Contingency Control mice when they were in the social conditioning chamber for pre-exposure or social conditioning. To encounter their assigned target, Pre-Contingency Control mice were placed into a clean cage with a clear acrylic divider separating them from their target. This divider allowed visual, olfactory, and auditory exchanges with the target but minimized tactile interactions, the same as for Social Stimulus + Stress mice and Shock Controls when in the social conditioning context. On Day 0, Pre-Contingency Control mice encountered their target for five min; on Day 1, for nine min; these were timed to match the length of target exposure that mice in the Social Stimulus + Stress group experienced. Post-Contingency Control (E) mice were treated the same as Pre-Contingency Control mice, except the former’s encounters with their assigned targets occurred four h *after*, rather than before, pre-exposure and social conditioning.

#### Apparatus and Software

On Days 0, 1, and 3, experimental mice were placed in Coulbourn Instruments chambers (17.8 cm D × 17.8 cm W × 30.5 cm H; Allentown, PA). Chambers comprised two opposing aluminum walls each adjoining two clear acrylic walls. Targets were put inside a transparent enclosure placed within a corner of the social conditioning chamber. The transparent enclosure included 23 holes (each 0.32 cm diameter) on each of the two sides facing the experimental mouse to enable olfactory cue exchanges. Previous work demonstrates that olfaction is the most important modality when recognizing and interacting with novel same-species targets (see review (Sterley and Bains, 2021)); tactile (whiskers) and auditory modalities appear to only be recruited when recognizing cage mates (de la Zerda et al., 2022) in male mice (females were not studied). Chambers and enclosures were cleaned with 70% ethanol before and after each session; chambers had visible illumination, and one clear acrylic wall was marked with a blue dotted pattern. These components contributed olfactory and visual cues to the context, in addition to the tactile cue of the stainless steel grid floor for the experimental mouse. Cameras mounted above the chambers were used to record movements, and freezing behavior (≥0.25 s) on Days 0, 1, and 3 was quantified using FreezeFrame software (v. 5.201, Actimetrics, Wilmette, IL). Freezing – ceasing all movement save breathing – is an innate behavior expressed by mice in response to a real or perceived threat, enabling threat assessment while minimizing detection (see review (Blanchard and Blanchard, 1988)). All videos were manually reviewed to ensure the software satisfactorily distinguished between freezing behavior and mere absence of locomotion.

On Day 2, experimental mice were placed in a novel room (distinct from the room used for Days 0, 1, and 3) containing open arenas (41.9 cm W × 41.9 cm D × 39.6 cm H; Coulbourn Instruments). Each open arena had one empty PVC tube (8.9 cm outer diameter) in a single corner of the square chamber with wire mesh (0.25 × 0.25 cm openings) covering a rectangular portion (11.4 cm W × 4.1 cm H) cut out of the tube bottom. Because of the mesh at the bottom of the PVC tubes containing targets, exchanges of visual, auditory, and olfactory cues between experimental mice and targets were possible, but tactile interactions were minimized. Social interaction behavior was quantified as post-test duration when at least 80% of an experimental mouse’s body was in the interaction zone, a 7 cm radius zone (∼223 cm^2^ – 240 cm^2^, i.e., 14-15% of the open arena) around the base of the PVC tube. Freezing behavior, regardless of arena location, lasting at least 250 ms during the post-test was also measured, then converted into a percentage of the post-test time. Social and freezing behaviors were detected using ANY-maze software (v. 7.09 Stoelting Co., Wood Dale, IL).

#### Pre-exposure (Day 0)

Pre-exposure was used to help reduce the novelty of the transparent enclosure and/or the target to the experimental mouse, plus minimize any potential sex differences in acquisition (Siegfried et al., 1987; Wiltgen et al., 2001; Guzmán et al., 2009; Brown et al., 2011). On Day 0, experimental mice were placed in chambers with (Shock Control and Social Stimulus + Stress mice) or without (Social, Pre-Contingency, and Post-Contingency Controls) a target (age-matched; either same sex or different sex relative to experimental mouse, depending on if assigned to same sex or different sex experiment) for pre-exposure (Fig. 1). During pre-exposure, experimental mice were allowed to explore the chamber and target enclosure for five min, then both mice were returned to their respective home cages. No aversive stimulus was presented.

#### Social Conditioning (Day 1)

On Day 1, experimental mice were placed in the same social conditioning chamber as Day 0 and received five, 1 s mild foot shocks (1.0 mA) in the presence of the same target that they encountered on Day 0 (Fig. 1). This was to have experimental mice form associations between aversive foot shocks and their respective target (Fig. 1). Conditioning lasted for nine min; after a two min baseline, shocks were then administered at 120, 210, 300, 390, and 480 s. Targets did not receive foot shocks.

#### Social Engagement Testing (Day 2)

On Day 2, experimental mice intentionally underwent social engagement testing in a novel environment with a novel target enclosure to ensure sensitivity of our social interaction measure while minimizing potential contextual confounds; in other words, to isolate effects on social engagement from those of context fear learning (the latter assessed on Day 3). Mice acclimated to the room in their home cage for 30 min prior to behavior testing commencing. Experimental mice were placed in the corner of the arena opposite the PVC tube and could investigate freely for 2.5 min (pre-test). Immediately after these 2.5 min, the target that each experimental mouse had previously been conditioned with on Day 1 was then placed in the PVC tube in the corner. Experimental mice were allowed to continue investigating the arena for an additional five min (post-test; Fig. 1). Proprietary incompatibilities and video file encoding precluded analyses of freezing behavior during social interaction testing from being evaluated with FreezeFrame software; this is why ANY-maze software was used instead here. Post-test social interaction time was log-transformed (Y=log(Y+0.001) to account for three mice with zero s post-test social interaction) so that data would be normally distributed for statistical analyses. Females were intentionally not assessed for estrus cycle stage for five reasons: 1) to minimize mouse usage (Russell and Burch, 1959), we did not power our studies for assessment of estrus; 2) with the goal of developing a social stress paradigm, we are seeking effects robust enough to not depend upon estrus in intact, randomly cycling females; 3) we intentionally focused here on fear and social behaviors, and did not measure sexual behaviors (e.g., lordosis); 4) evidence that overall mouse behaviors are not affected by estrus stage (Prendergast et al., 2014; Levy et al., 2023), (but see (Chari et al., 2020)); 5) cross-species evidence indicates vaginal lavage to determine estrus cycle is stressful (Becegato et al., 2021; Bahadır-Varol et al., 2022), and we sought to minimize stress confounds here.

#### Context Testing (Day 3)

On the last day of the paradigm, experimental mice were placed in the social conditioning chamber for behavior testing in the *absence* of any target (Fig. 1). The target enclosure was still present to keep the context consistent. All other tactile, visual, and olfactory cues from Days 0 and 1 were present. Testing lasted 10 min and did not involve any foot shocks.

### Treatment Groups

The Social Stimulus + Stress group involved mice that underwent social conditioning in the presence of a non-shocked target (either same sex or different sex relative to Social Stimulus + Stress mouse, depending upon experiment), with the hypothesis that Social Stimulus + Stress mice would associate the target with this aversive experience. We planned four Control groups for this one Social Stimulus + Stress group; each experiment (same sex or different sex) had its own respective set of four Control groups for its respective Social Stimulus + Stress group. The goals of these were to control for: 1) foot shock exposure, i.e., Shock Controls; 2) presence of target during pre-exposure & foot shock, i.e., Social Controls; 3) exposure to target temporally distal to foot shock exposure, such that social encounters still occurred in a manner not contingent with foot shock nor the conditioning context, i.e., Pre– and Post-Contingency Controls (Fig. 1). Shock Controls experienced procedures identical to those of the Social Stimulus + Stress group, except a foot shock was never administered on Day 1 (Fig. 1). Social Controls were treated the same as the Social Stimulus + Stress group, except experimental mice never encountered their assigned target until testing on Day 2 (Fig. 1). Pre– and Post-Contingency Controls involved experimental mice encountering targets either four h before (Pre-Contingency Controls) or four h after (Post-Contingency Controls) pre-exposure (Day 0) and social conditioning (Day 1); in other words, targets were not present in the conditioning chamber during these two periods. Instead, experimental mice were exposed to their target in a separate room for five min or nine min (the same amount of time as pre-exposure or social conditioning, respectively) four h before (Pre-Contingency Controls) or four h after (Post-Contingency Controls) the experimental mice were exposed to the conditioning chamber. This exposure involved experimental mice and their respective target being placed in a clean mouse cage separated with an acrylic divider to allow for visual, auditory, and olfactory exchange, but minimizing tactile interaction, mirroring the experiences of Social Stimulus + Stress mice and Shock Controls within the social conditioning context on Days 0-1. Mice in Pre-or Post-Contingency Control conditions still experienced foot shocks on Day 1, and encountered their assigned target for testing on Day 2 (Fig. 1). This temporal separation of four h was to minimize consolidation of social encounters from interfering/intermingling with consolidation of the aversive foot shock encounter in the social conditioning chamber, while still making execution of these experiments feasible within a 12 h lights-on period (Wanisch et al., 2008; Cai et al., 2016; Lynch et al., 2017) (see reviews (Abel and Lattal, 2001; Sheppard et al., 2018)).

### Serum Corticosterone

Thirty min after all experimental mice underwent contextual fear testing on Day 3, they were briefly anesthetized with isoflurane then rapidly decapitated for trunk blood collection. Blood clotted at room temperature for 10 min, then was spun at 3500 rpm for one h at 4°C. Serum was collected and stored at –80°C until corticosterone analyses could be performed. Serum corticosterone was measured using Enzo Life Sciences corticosterone enzyme-linked immunosorbent assay kits (Farmingdale, NY). Assays were run according to the manufacturer’s instructions using their small volume protocol. Plates were read at 405 nm with correction at 580 nm. The sensitivity of the assay was 26.99 pg/mL. After serum corticosterone levels were interpolated from each plate’s standard curve, they were log-transformed to account for the typical skewness of serum corticosterone (Teilmann et al., 2014; Uarquin et al., 2016; Weber et al., 2023).

### Statistical Analyses

Data were graphed with GraphPad Prism 10.3.0 (442), Beta (GraphPad Software, San Diego, CA), and analyzed using IBM SPSS Statistics 28.0.0.0 (IBM, Armonk, NY), with the significance threshold set *a priori* at p<0.05. Non-significant trends (p<0.10) were mentioned only when the associated partial η**^2^**was ≥0.060. Data were graphed as the mean (μ) ± standard error of the mean. Details of identified outliers (greater than the mean ± four standard deviations (σ)) are provided in the Supplemental Dataset. The same sex experiment and different sex experiment each included their own Social Stimulus + Stress group plus accompanying four Control groups. Experiments were analyzed separately. Social conditioning acquisition was analyzed using a three-way repeated measures general linear model (time × treatment group × sex of experimental mouse). Greenhouse Geisser corrections were utilized for within-subjects analyses. Measurements of contextual fear expression average, log-transformed post-test social interaction time, percent time freezing during social interaction testing, and log-transformed corticosterone were each analyzed with a two-way general linear model (treatment group × sex of experimental mouse). Our primary outcomes of interest for this study were freezing during social conditioning acquisition, social engagement behavior, and freezing during context testing. Accordingly, our *a priori* hypotheses and planned contrasts were:

- **Hypothesis 1**: Freezing during the final time point of social conditioning acquisition would be minimal in Shock Control mice compared to all other groups; and female experimental mice would exhibit higher levels of freezing than males.
- **Hypothesis 2**: In female more than male experimental mice, log-transformed social engagement would be decreased in Social Stimulus + Stress mice compared to all four Control groups.
- **Hypothesis 3**: Freezing during testing in the social conditioning chamber would be minimal in Shock Control mice compared to all other groups; and female experimental mice would exhibit higher levels of freezing than males.

These planned contrasts were selectively analyzed without correction alongside our omnibus analyses. For our secondary outcomes that either were used to assess the possibility of confounds (percent time freezing during social interaction testing) or were exploratory (log-transformed serum corticosterone), and as follow-ups to our planned contrasts on our primary outcomes, we performed *post hoc* pairwise comparisons with Bonferroni correction. The results of these *post hoc* tests for our primary outcomes are reported in Supplemental Figures S1-S3.

## Results

### Social Conditioning Acquisition

Social conditioning acquisition is illustrated in Fig. 2A, 2B for mice in the same sex target experiment, and in Fig. 2C, 2D for mice in the different sex target experiment. All mice receiving a foot shock expressed >40% freezing during at least one post-shock period during social conditioning.

**Figure 2.**
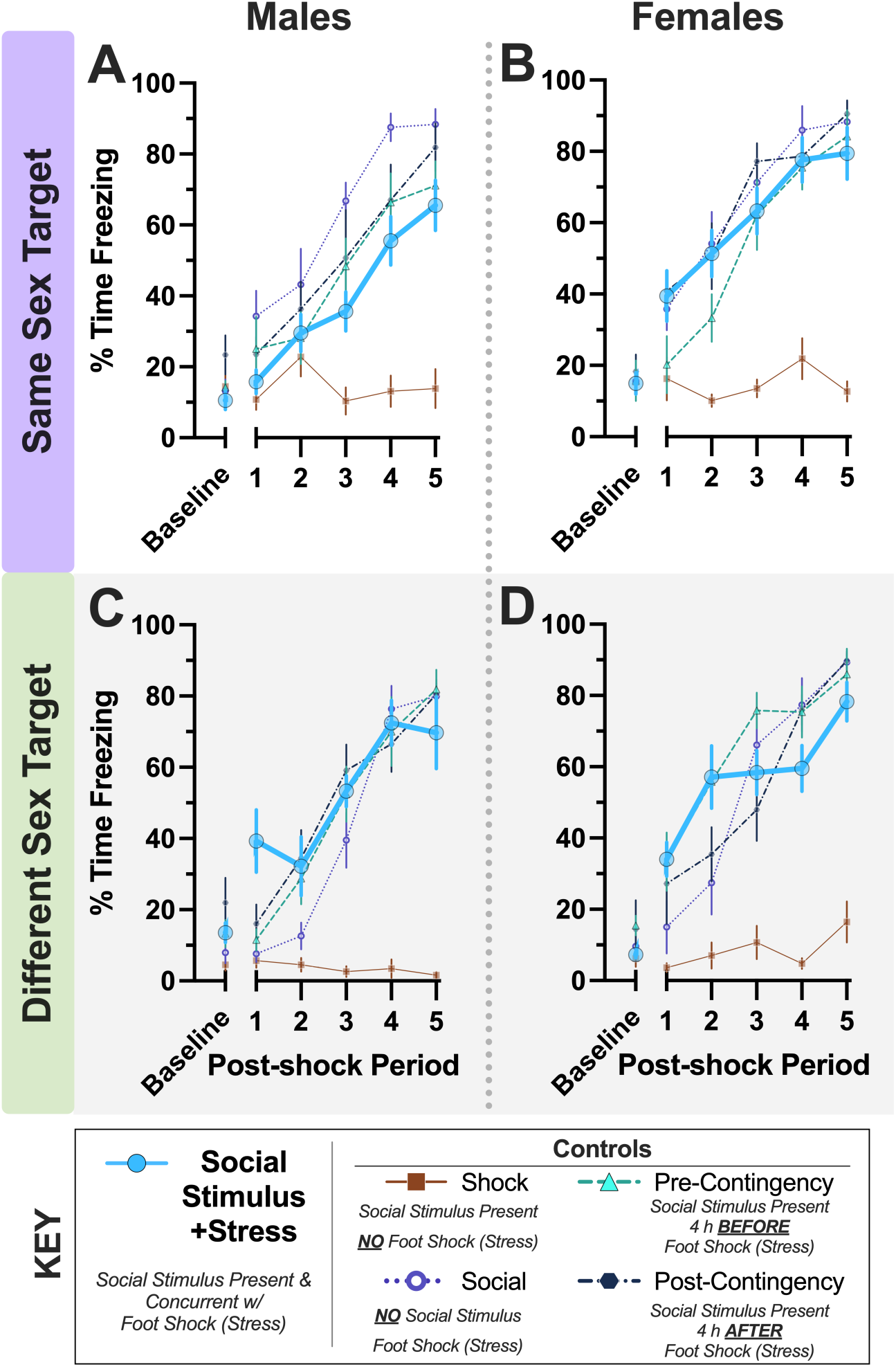
Day 1 acquisition during social conditioning procedure. Percent time freezing during social conditioning acquisition for mice in same sex (Panels A, B) and different sex (Panels C, D) experiments for male (A, C) and female (B, D) mice. Numbers of mice graphed within Panels A-D in order: Social Stimulus + Stress (n=8, 9, 9, 8); Shock Control (8, 8, 7, 7); Social Control (n=8 for all); Pre-Contingency Control (n=7, 8, 8, 9); Post-Contingency Control (n=7, 8, 8, 8). Average freezing for the first two min, prior to commencement of acquisition, is plotted on the x-axis as baseline. The 410 average percent freezing for each 30 second period following each of the five mild foot shocks are thereafter plotted along x-axis (Post-shock Periods 1-5). Data graphed as mean ± standard error of the mean.

For acquisition in the same sex experiment, no three-way interaction of time × sex × group occurred (Table 1). We observed a significant two-way interaction between time × group, and a non-significant trend for group × sex was noted (p=0.070; Table 1). A significant main effect of sex was also found for mice in the same sex experiment (Table 1). Our *a priori* planned contrast for **Hypothesis 1** within the same sex experiment (F(_1,69_)=1.140 p=0.289, partial η^2^=0.016) did not support our anticipated observation (Fig. 2A, B). Graphs of these data showing the outcomes of pairwise comparisons are located in Supplemental Figure S1A, B.

**Table 1.**
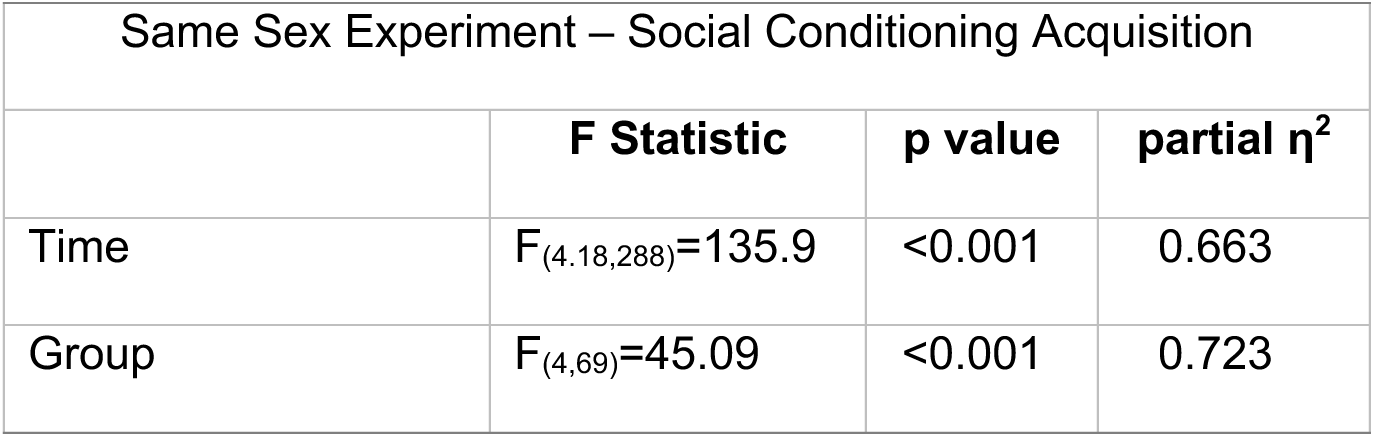
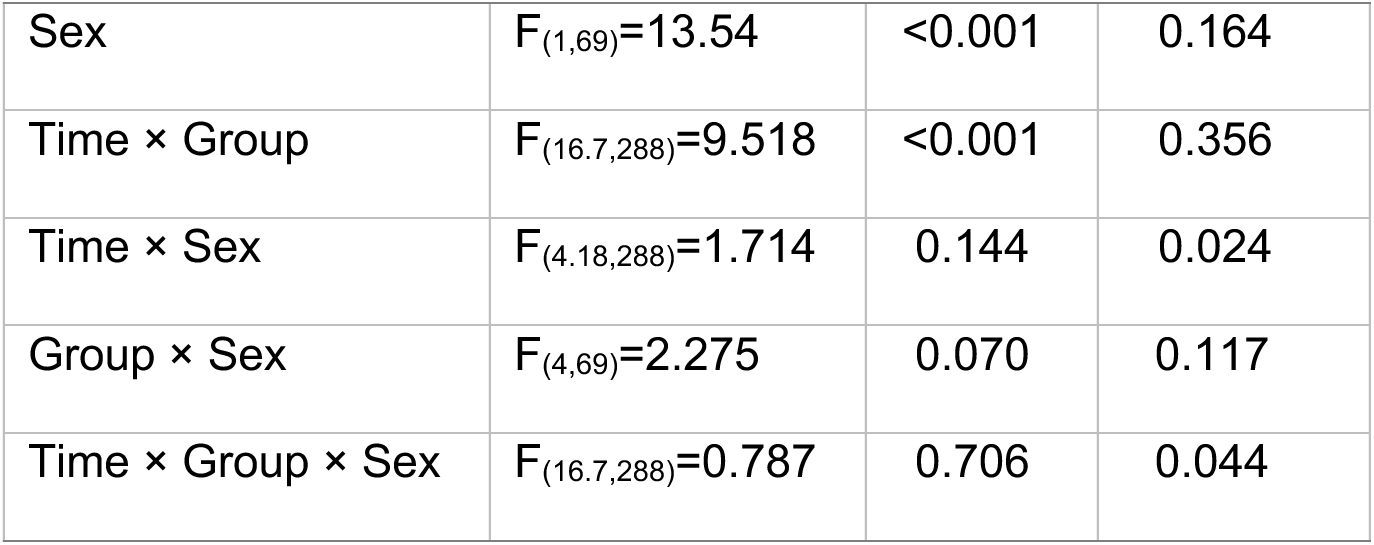
Two-way repeated measures general linear model on learning of social conditioning for mice of both sexes with same sex targets.

Social conditioning acquisition for mice in the different sex experiment is shown in Fig. 2C, 2D. A significant three-way interaction of time × sex × group was observed (Table 2). Similar to the same sex experiment, however, our planned contrast did not reach significance for **Hypothesis 1** (F(_1,70_)=0.537 p=0.466, partial η^2^=0.008). Indeed, no significant pairwise comparisons indicated within-group significant sex differences at the final social conditioning acquisition time point, either for the same sex or different sex experiments (Supplemental Figure S1).

**Table 2.** Two-way repeated measures general linear model on learning of social conditioning for mice of both sexes with different sex targets.

| Different Sex Experiment – Learning Social Conditioning |  |  |  |
| --- | --- | --- | --- |
|  | **F Statistic** | **p value** | **partial $\eta^2$** |
| Time | $F_{(4.35,304)}=184.2$ | <0.001 | 0.725 |
| Group | $F_{(4,70)}=38.88$ | <0.001 | 0.690 |
| Sex | $F_{(1,70)}=6.694$ | 0.012 | 0.087 |
| Time × Group | $F_{(17.4,304)}=12.67$ | <0.001 | 0.420 |
| Time × Sex | $F_{(4.35,304)}=3.317$ | 0.009 | 0.045 |
| Group × Sex | $F_{(4,70)}=0.892$ | 0.474 | 0.048 |
| Time × Group × Sex | $F_{(17.4,304)}=1.658$ | 0.048 | 0.087 |

Combined with same sex experiment acquisition (Supplemental Figure S1), these findings indicate that social exposure – whether concurrent with or temporally distal from an aversive unconditioned stimulus – can have transient impacts on the freezing behavior exhibited by experimental mice across sexes. All conditioning procedures here, though, culminated in similar final freezing levels across sexes of both experimental mice and their targets. This facilitates comparisons of the different conditioning manipulations on subsequent social and freezing behaviors by minimizing potential acquisition confounds.

### Social Interaction Testing

Social conditioning in the same sex experiment resulted in a sex × group interaction on social engagement (Table 3). Moreover, our planned contrast supported **Hypothesis 2** for the same sex experiment (Fig. 3A; F(_1,68_)=7.037 p=0.010, partial η^2^=0.094), though our contrast estimate was negative. This informed us that our proposed directionality of the sex × group interaction was not supported by the data. In evaluating means for our planned contrast, we found that log-transformed social engagement values for Social Stimulus + Stress males (μ=1.008, σ=1.698, n=8) were indeed decreased compared to all male Controls (μ=2.036, σ=0.269, n=29). However, the mean for Social Stimulus + Stress females (μ=1.813, σ=0.991, n=32) was actually higher than that for all pooled female Controls (μ=1.968, σ=0.417, n=9), likely due to some Social Control females with low social engagement values (Fig. 3A). Pairwise comparisons (Supplemental Figure S2A, B) corroborate the low mean for Social Control females, plus indicate that Social Stimulus + Stress males displayed less social engagement than Social Stimulus + Stress females, reiterating that our paradigm was not successful in its goal.

**Figure 3.**
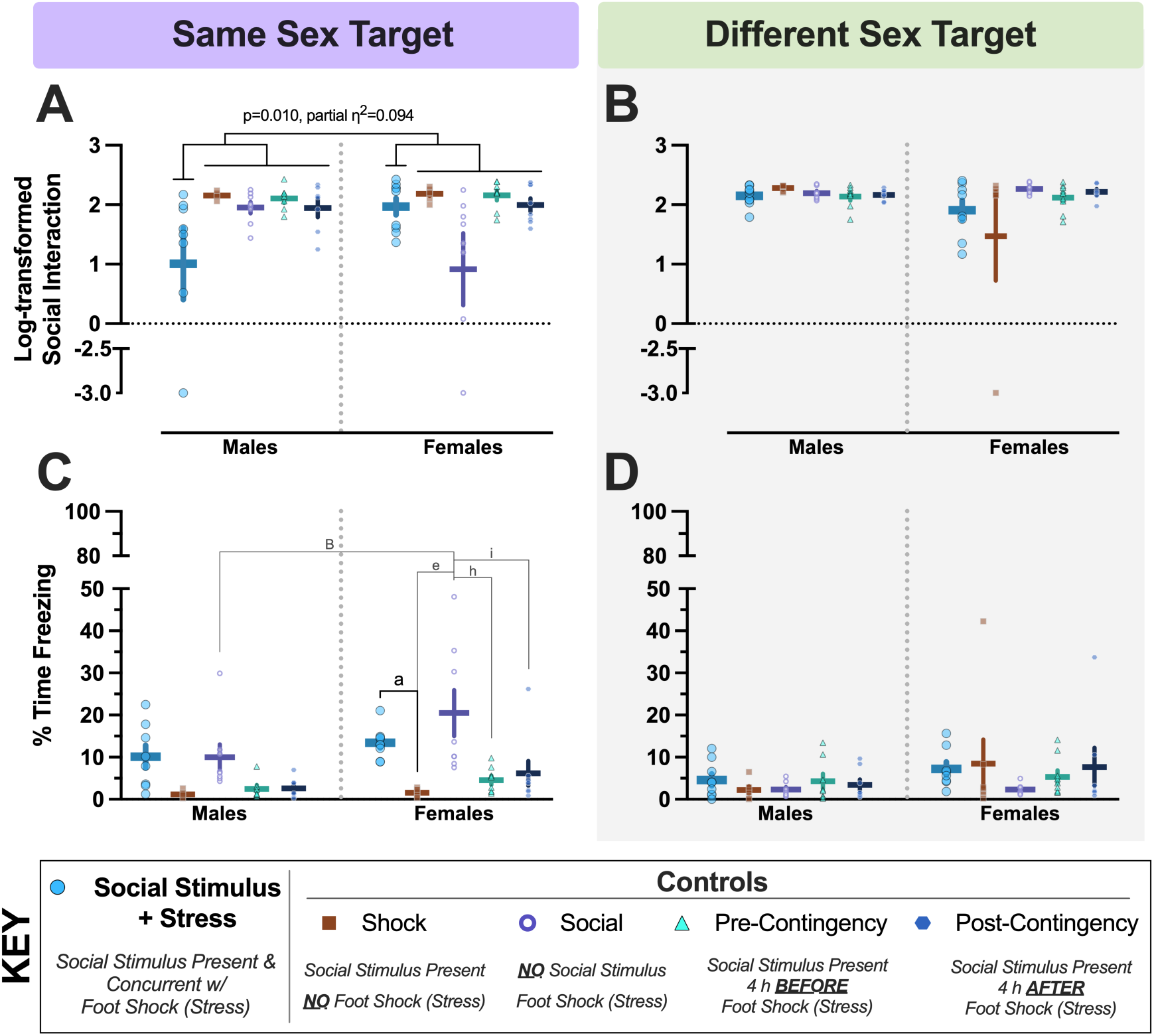
Day 2 social interaction and fear behaviors following social conditioning. Log transformed social interaction (Panels A, B) and percent time freezing during post-test social interaction (Panels C, D) data are shown for mice in the same (A, C) and different (B, D) sex experiments. Numbers of mice graphed within each panel, left to right: A) n=8, 7, 8, 7, 7, 9, 8, 8, 8, 8; B) n=9, 6, 8, 8, 8, 8, 7, 7, 9, 7; C) n=8, 7, 8, 7, 7, 9, 8, 8, 8, 8; D) n=9, 6, 8, 8, 8, 8, 7, 7, 9, 7. Y axes for A, B were split to visualize data points for three total mice across three separate groups that exhibited no social interaction. Y axes for C, D were split to facilitate clearer visualization of the low freezing levels exhibited during social engagement testing. An a priori planned contrast evaluating if log-transformed social interaction was reduced in Social Stimulus + Stress mice across sexes compared to all four Control groups within the same experiment was significant for mice in the same sex experiment (Panel A; F_(1,68)_=7.037 p=0.010, partial η^2^=0.094). Results of pairwise comparisons with Bonferroni correction, left to right in Panel C: ^B^indicates (Social Controls male vs. female) p=0.004; ^a^indicates (Shock Control vs. Social Stimulus + Stress) p=0.008; ^e^indicates (Shock Control vs. Social Control) p<0.001; ^h^indicates (Social Control vs. Pre-Contingency Control) p<0.001; ^i^indicates (Social Control vs. Pre-Contingency Control) p=0.001. Data graphed as mean ± standard error of the mean.

**Table 3.** Two-way general linear model on social interaction for mice of both sexes with same sex targets.

| Same Sex Experiment – Log-transformed Social Interaction |  |  |  |
| --- | --- | --- | --- |
|  | **F Statistic** | **p value** | **partial $\eta^2$** |
| Group | $F_{(4,68)}=2.967$ | 0.025 | 0.149 |
| Sex | $F_{(1,68)}=0.004$ | 0.951 | 0.000 |
| Group × Sex | $F_{(4,68)}=3.097$ | 0.021 | 0.154 |

Fear behavior, indexed as percent time freezing during the social interaction post-test, was also evaluated. While the goal of our modified paradigm was to ultimately decrease social engagement following conditioning, we also assessed the level of freezing expressed in the novel social testing arena. In the same sex experiment, no sex × group interaction occurred, but main effects of both group and sex were observed (Table 4). Pairwise comparisons indicated that freezing was higher in Social Stimulus + Stress females versus Shock Control females, while the same comparison in males revealed no differences (Fig. 3C). In a somewhat inverted pattern compared to Controls during same sex social interaction, Social Control females exhibited more freezing during social interaction testing than Social Control males, Shock Control females, and Pre– and Post-Contingency Control females (Fig. 3C). Overall, though, freezing levels during social interaction testing were minimal as compared to those occurring during acquisition (Fig. 2) and context testing (Fig. 4A-B).

**Figure 4.**
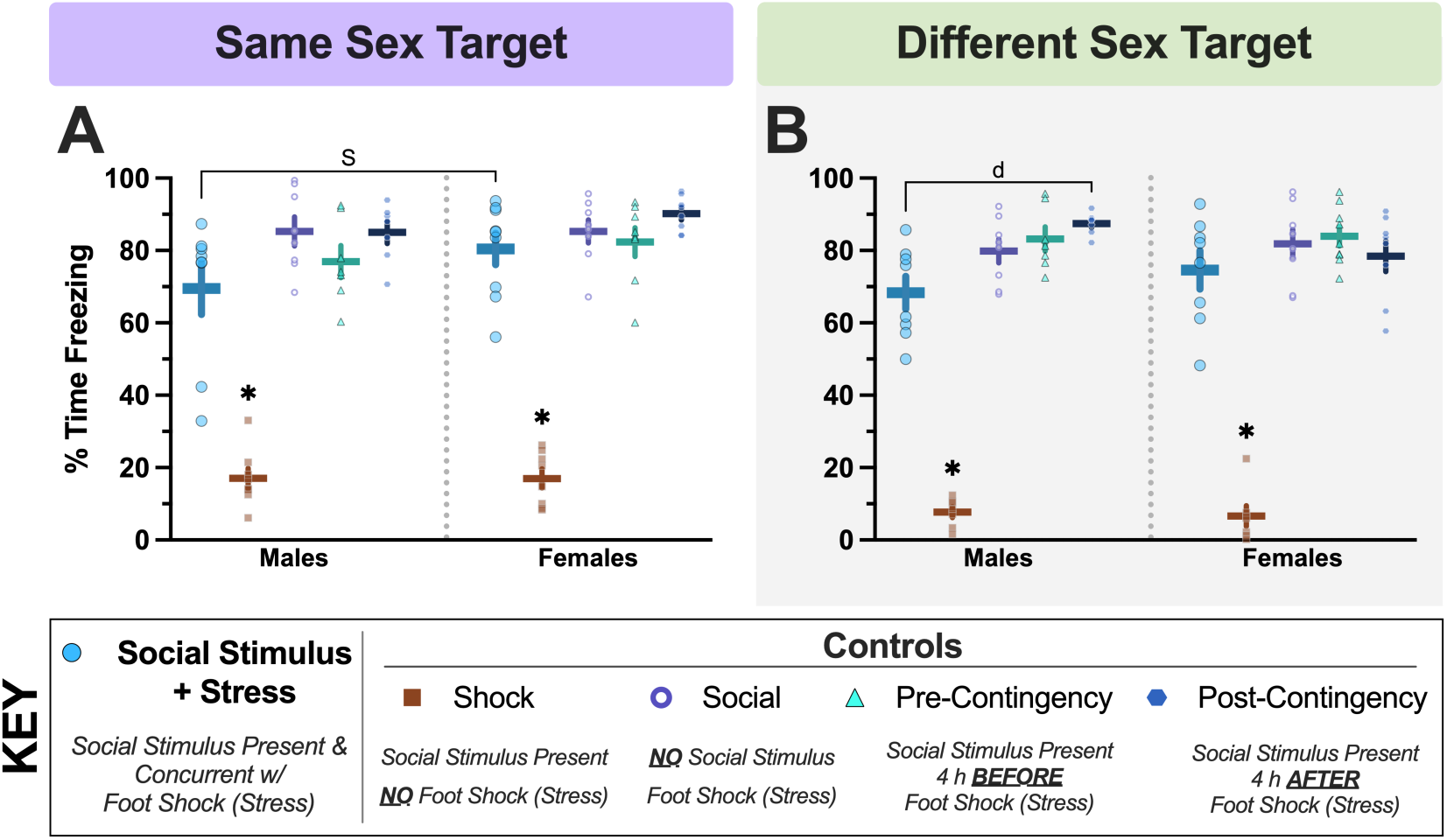
Day 3 social conditioning context fear expression averages. Average context fear behavior (Panels A, B) data are shown for mice in all groups in the same sex (A) and different sex (B) experiments. Numbers of mice graphed within Panels A-B in order, left to right: A) n=8, 8, 8, 7, 7, 9, 8, 8, 8, 8; B) n=8, 7, 8, 7, 7, 8, 7, 8, 9, 8. Left to right: ^S^p=0.049, ^d^indicates (Pre-Contingency Control vs. Social Stimulus + Stress) p=0.004. ✱p=0.001 Social, Pre-Contingency, and Post-Contingency Controls and Social Stimulus + Stress mice vs. Shock Control. Data graphed as mean ± standard error of the mean.

**Table 4.**
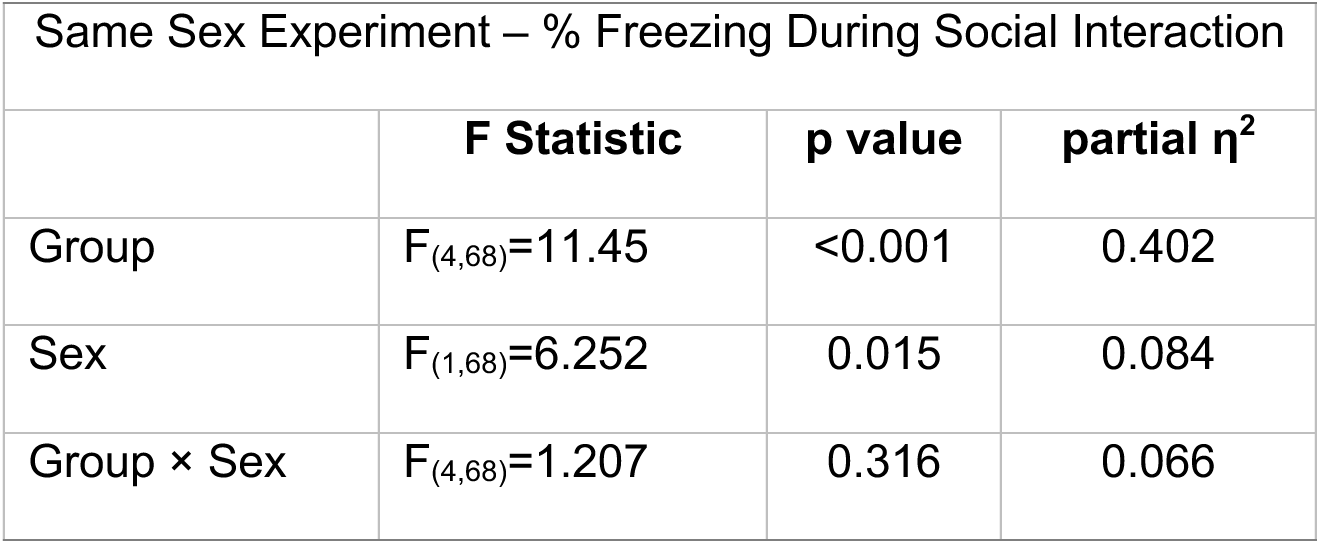
Two-way general linear model on percent time freezing during social interaction for mice of both sexes with same sex targets.

Contrasting with the same sex experiment, there was no significant interaction nor any main effects in the different sex experiment for social engagement (Table 5). Further distinguishing the different sex experiment with the same sex experiment, our **Hypothesis 2** planned contrast for the different sex experiment did not achieve significance (Fig. 3B; F(_1,67_)=0.033 p=0.856, partial η^2^=0.000). Pairwise comparisons demonstrated that Social Stimulus + Stress did not generate any sex differences in social engagement (Supplemental Figure S2B). The only sex effect observed was in Shock Controls, with females demonstrating less social engagement than males, though this appears driven by a single Shock Control female (Supplemental Figure S2B). Combined, these findings indicate that the social engagement of female mice with other females is more affected by prior aversive experiences in the *absence* of any social stimuli, whereas males are more socially affected by temporally concurrent aversive associations with their own sex.

**Table 5.** Two-way general linear model on social interaction for mice of both sexes with different sex targets.

| Different Sex Experiment – Log-transformed Social Interaction |  |  |  |
| --- | --- | --- | --- |
|  | **F Statistic** | **p value** | **partial $\eta^2$** |
| Group | $F_{(4,67)}=0.726$ | 0.577 | 0.042 |
| Sex | $F_{(1,67)}=1.756$ | 0.190 | 0.026 |
| Group × Sex | $F_{(4,67)}=1.172$ | 0.331 | 0.065 |

Unlike the same sex experiment, neither an interaction nor main effects of sex and group were observed with freezing behavior during social interaction testing for the different sex experiment (Table 6). Moreover, no significant pairwise comparisons were observed (Fig. 3D). The predominant consistency across experiments for freezing was that expression of fear was minimal during the post-test of social interaction, as compared to freezing quantified in the conditioning context (Figs. 2, 4A-B).

**Table 6.** Two-way general linear model on percent time freezing during social interaction for mice of both sexes with different sex targets.

| Different Sex Experiment – % Freezing During Social Interaction |  |  |  |
| --- | --- | --- | --- |
|  | **F Statistic** | **p value** | **partial $\eta^2$** |
| Group | $F_{(4,67)}=0.698$ | 0.596 | 0.040 |
| Sex | $F_{(1,67)}=3.414$ | 0.069 | 0.048 |
| Group × Sex | $F_{(4,67)}=0.491$ | 0.743 | 0.028 |

### Social Conditioning Context Fear Testing

Because of the natural extinction process that can occur during testing, we specifically examined behavioral expression of social conditioning context fear during minutes two through six of testing, to best capture fear expression with minimal confounds from extinction processes (Lynch et al., 2017; Beaver et al., 2023; Weber et al., 2023). This testing further allowed us to determine if social engagement testing on Day 2 might have any carry-over effects on contextual fear expression on Day 3 (Tables 7-8, Fig. 4A-B).

**Table 7.** Two-way general linear model on both sexes’ context fear testing average with same sex targets.

| Same Sex Experiment – Fear Expression Average |  |  |  |
| --- | --- | --- | --- |
|  | **F Statistic** | **p value** | **partial $\eta^2$** |
| Group | $F_{(4,69)}=108.5$ | <0.001 | 0.863 |
| Sex | $F_{(1,69)}=2.859$ | 0.095 | 0.040 |
| Group × Sex | $F_{(4,69)}=0.676$ | 0.611 | 0.038 |

**Table 8.**
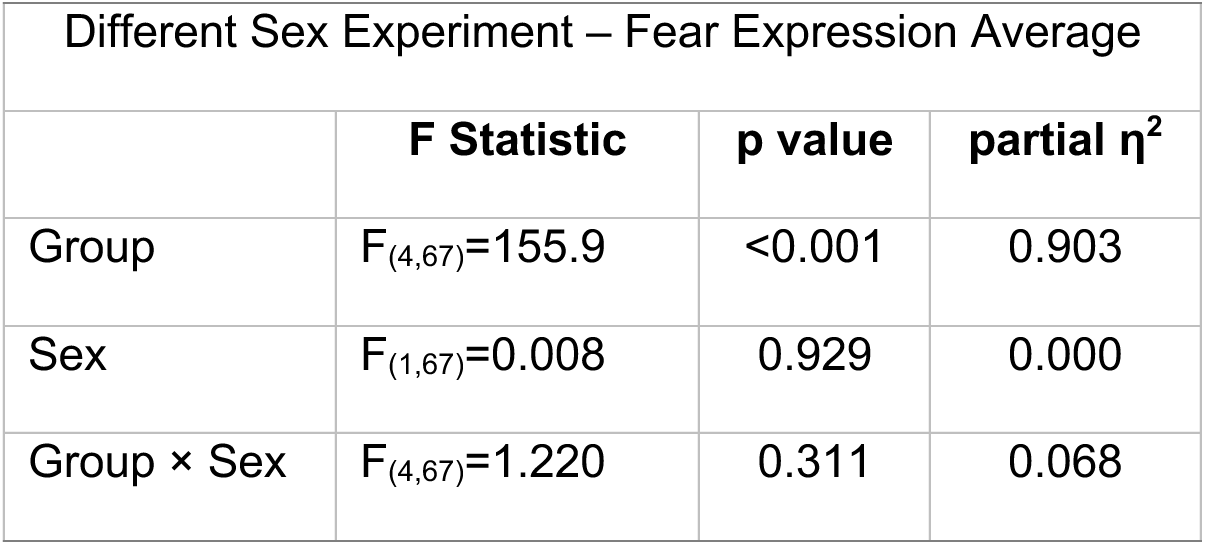
Two-way general linear model on both sexes’ context fear testing average with different sex targets.

For the same sex experiment, no two-way interaction between group × sex was found, but a significant main effect of group emerged (Table 7). Given omnibus statistics did not report an interaction, it is not surprising that our planned contrast for **Hypothesis 3** in the same sex experiment likewise was not significant (Fig. 4A; F(_1,69_)=0.733 p=0.395, partial η^2^=0.011). This was further supported by pairwise comparisons (Supplemental Figure S3A), which indicated that male Social Stimulus + Stress mice exhibited significantly less average freezing in the conditioning context compared to female Social Stimulus + Stress mice (Fig. 4A). Shock Control mice of both sexes exhibited reduced fear behavior compared to all other groups, as expected (Fig. 4A).

In the different sex experiment, no significant group × sex interaction was found, nor was a significant effect of sex (Table 8). A significant group effect was similarly observed for the different sex experiment (Fig. 4B). As with the same sex experiment, the lack of a significant omnibus interaction was reflected in the non-significant outcome of our planned contrast for **Hypothesis 3** (F(_1,67_)=0.037 p=0.848, partial η^2^=0.001). Pairwise comparisons once more supported low freezing in Shock Controls across sex relative to all other groups, plus indicated that male Social Stimulus + Stress mice froze significantly less than Post-Contingency Control males (Supplemental Figure S3B).

Integrated with the results from social interaction testing, these findings indicate that conditioned shifts in social engagement are distinct from conditioned fear to the social conditioning context. In parallel, freezing behavior was relatively minimal during social interaction testing compared to testing in the conditioning context, suggesting both that freezing behavior did not confound social interaction outcomes and that mice were able to sufficiently discriminate between contexts. Curiously, interaction of male Social Stimulus + Stress mice with familiar targets in a novel environment on Day 2 might lead to mild reductions in contextual fear expression on Day 3.

### Serum Corticosterone

Log-transformed corticosterone levels of mice in the same sex experiment exhibited no significant interaction of group × sex, but a main effect of group was observed (Supplemental Table S1, Supplemental Figure S3C). Pairwise comparisons showed that, across sex, Social Stimulus + Stress mice in the same sex experiment exhibited elevated log-transformed corticosterone levels compared to their respective same sex Shock Control mice (Supplemental Figure S3C). While this was not surprising, male Pre– and Post-Contingency Control mice also exhibited higher log-transformed corticosterone levels than Shock Control mice, whereas Social Control mice did not (Supplemental Figure S3C). Conversely, female Social Control mice had higher log-transformed corticosterone levels than Shock Control mice, but female Pre– and Post-Contingency Control mice did not (Supplemental Figure S3C). Given no target was present during Day 3 testing, these findings suggest enduring and sex-specific physiological differences following social conditioning, despite minimal behavioral shifts. Specifically, any social interaction with their assigned target on Day 1, whether concurrent or temporally separate with foot shock exposure, elevated male log-transformed corticosterone levels but attenuated female log-transformed corticosterone levels after context testing on Day 3 *in the absence of any targets*. Only Social Stimulus + Stress females did not align with this pattern, exhibiting heightened log-transformed corticosterone levels on Day 3 similar to those of Social Control females (Supplemental Figure S3C).

Log-transformed corticosterone levels of mice in the different sex experiment, while not exhibiting a significant sex × group interaction, did display significant main effects of both group and sex (Supplemental Table S2). In contrast to the same sex experiment, pairwise comparisons indicated that within each sex, levels of log-transformed corticosterone did not differ across groups, including relative to Shock Control mice (Supplemental Figure S3D). The only difference that reached significance was male Social Stimulus + Stress mice having lower log-transformed corticosterone levels compared to female Social Stimulus + Stress mice (Supplemental Figure S3D). Combined with log-transformed corticosterone data from the same sex experiment (Supplemental Figure S3C), these findings illustrate how same sex target interactions on the same day as an aversive learning experience can elicit sex-specific and enduring hormone changes. Moreover, these lasting physiological shifts are not mirrored after different sex target interactions. This indicates the combination of target and experimental sex, when temporally concurrent or adjacent to a fear learning event, influences endocrine responses to context days later.

## Discussion

Our goal in executing these experiments was to begin establishing a paradigm for inducing consistent, reproducible socially paired stress in mice across sexes. While our results indicate this specific paradigm does not yet accomplish our goal, we want to share our approach and findings so that others can utilize this information to efficiently prioritize future efforts in this domain. Moreover, despite a mostly negative outcome towards our overarching goal, we nevertheless uncovered sex-specific patterns of behavioral and physiological stress responsivity that are useful to fields incorporating sex as a biological variable, and/or seeking to employ different sex social stressors in mice.

All mice that underwent social conditioning with mild foot shocks (Social, Pre-Contingency, and Post-Contingency Controls plus Social Stimulus + Stress mice) ultimately ended their acquisition stage at similar freezing levels. Subsequent outcomes in behavioral testing for social engagement and contextual fear expression, as well as for serum corticosterone, could, therefore, be interpreted with minimal concern for acquisition confounds. Conversely, this consistency might have contributed to our negative findings. Alternatively, they might indicate that our social conditioning protocol resulted in a ceiling effect. Additional studies would need to parse this out.

Our social interaction findings were not as robust as anticipated. For our same sex experiment, we found that male Social Stimulus + Stress mice engaged less during social testing compared to Shock Control mice, and compared to Social Control mice that never encountered a target during training. Additionally, male Social Stimulus + Stress mice exhibited less social engagement than female Social Stimulus + Stress mice in same sex experiment, whereas social engagement was unaffected across sexes in the different sex experiment. This reveals that male mice likely form stronger associations when aversive experiences occur in conjunction with the presence of another male mouse, whereas female mice appear to reduce social engagement after experiencing non-social aversive stimuli. Such a sex difference could help explain some of the challenges in developing social stress paradigms using female laboratory mice (*Mus musculus*). Relatedly, this could indicate that, in the presence of a potential sexual partner, threat assessment is suppressed. The inverse has been demonstrated in female rats; that is, fear attenuates sexual behaviors, at least in part through amygdala and hypothalamus estrogen receptor signaling (Moëne et al., 2019). Indirect evidence in male rats housed with females post-context fear conditioning suggests that encounters with different sexes may suppress threat assessment. These male rats subsequently exhibited attenuated fear expression, involving dopamine receptor signaling in the hippocampus (Bai et al., 2009).

However, a different group studying mice reported that ejaculation by males was required for retention of extinguished social fear conditioning (Grossmann et al., 2024). Continued evaluations are needed to determine whether our observations – under conditions where copulation is impossible – that male exposure to a female can diminish threat assessment behaviors are reproducible. Evidence for social transmission of stress in rodents suggests an alternative interpretation (Brandl et al., 2022). Instead, it could be that because the target for each Social Stimulus + Stress mouse did not experience the same stress (mild foot shock), this in turn affects the Social Stimulus + Stress mouse’s perception of their own aversive experience (Guzmán et al., 2009), likely in a sex-specific manner.

These sex-specific behavior patterns correspond to evidence that accumbal dopamine signaling dynamics in mice are sex-dependent, both upon the sex of the studied mouse and the sex of their assigned target (Dai et al., 2022). Our findings align with the success of using social defeat to stress male mice. Additionally, they indicate that future studies seeking to suppress female mice’s social interaction behavior through a socially associated stimulus will meet with more success if using an approach distinct from ours, which produced null results. Indeed, others have encountered challenges when trying to elicit behavioral shifts in female mice, even after four weeks of stress that involved only female targets (Díez-Solinska et al., 2023). One group reported that female mice consistently prefer a socially paired food reinforcer, even if the pairing was with a same sex target that had just previously undergone acute stress (mild foot shock) (Kietzman et al., 2022). Because males’ food preference was not studied by the researchers, and neither did their females encounter stressed different sex targets, it remains unclear if these effects are sex-specific, and if they would generalize to different sex interactions. Future studies will also need to determine how socially conditioning female mice to male targets subsequently affects interactions with female targets, and vice versa.

One day after social engagement testing, mice were tested for fear expression in the social conditioning context in the absence of any targets. A sex difference in context fear expression was observed between Social Stimulus + Stress mice in the same sex experiment, but not in the different sex experiment. This is the only directionally comparable outcome with social engagement, in that male Social Stimulus + Stress mice in the same sex experiment both exhibited less context fear and less social engagement versus female Social Stimulus + Stress mice. This could be interpreted in at least three ways. Male Social Stimulus + Stress mice might have found their male target more salient than the context, thereby leading them to form stronger associations between the mild foot shocks and their targets as compared to female Social Stimulus + Stress mice. Another possibility is that when mice are in the vicinity of a possible sexual partner, they could experience comparably impaired encoding of context cues, though as discussed in the preceding three paragraphs, literature investigating such target-sex based encoding effects is limited. Alternatively, the presence of the male target may have led to social buffering of the mild foot shocks for male Social Stimulus + Stress mice (Guzmán et al., 2009; Brandl et al., 2022), whereas this process did not occur for female Social Stimulus + Stress mice in the same sex experiment. Because our goal was for mice to associate the aversive mild foot shock with their assigned target, any social buffering of social conditioning context fear could counteract our goal. This is why our use of targets of a different strain, that were never housed in the same room as experimental mice, ensured they had never previously been encountered (even via olfaction) by the mice tested here and therefore at baseline there was no familiarity. Given the critical contribution of attachment to social buffering (see review (Kiyokawa and Hennessy, 2018)), we can be reasonably confident that there were no social buffering effects upon social conditioning context fear in same sex experiment mice. Thus, we are inclined toward the first interpretation.

Serum was collected 30 minutes after context fear testing, 24 h after experimental mice’s last encounter with their assigned target. We were interested to discover that sex-specific group differences in log-transformed corticosterone levels in the same sex experiment appeared to be best explained by Day 1 group conditions, rather than behavior on Days 1-3. For example, male Shock Control mice’s log-transformed corticosterone levels were lower than Pre– and Post-Contingency Controls and Social Stimulus + Stress groups – the three conditions where both target exposure and foot shocks occurred within an eight h window on Day 1. Social Control male mice only experienced foot shocks, and did not encounter a target, on Day 1.

Shock Control males similarly only encountered targets, and did not experience foot shocks on Day 1; and male Social Control log-transformed corticosterone levels did not differ from Shock Control males. Like males in the same sex experiment, female Social Stimulus + Stress mice had elevated log-transformed corticosterone relative to Shock Control females. In contrast, however, with this experiment’s females, only Social Control females had higher log-transformed corticosterone levels than Shock Control females. Pre– and Post-Contingency Control females’ log-transformed corticosterone was not different from Shock Controls, and the former groups encountered targets four h before or after, respectively, experiencing foot shocks. Collectively, this suggests social encounters temporally distal from the foot shock, but still occurring within the same day, might augment males’ and mitigate females’ log-transformed corticosterone responses to a stressful experience *two days later*. In contrast, coinciding foot shocks and target exposure in Social Stimulus + Stress mice obscured this sex difference in endocrine response.

Unlike the same sex experiment, the different sex experiment mice’s log-transformed corticosterone levels did not map onto groups in either sex, despite a significant group effect being detected. Also distinct from the same sex experiment is that log-transformed corticosterone levels of Social Stimulus + Stress mice in the different sex experiment had a sex difference. Specifically, female Social Stimulus + Stress mice had greater log-transformed corticosterone levels than male Social Stimulus + Stress mice. This does not map onto their social engagement on Day 2, nor their average context fear expression on Day 3. This log-transformed corticosterone level difference could, therefore, be an enduring endocrine effect of encountering their assigned targets on Day 2.

To ensure we rigorously evaluated this modified paradigm’s hypothesized utility as a higher throughput approach for more consistent induction of social stress across sexes, we included four separate controls. Had we only included one Control group, e.g., Shock Controls, we might have incorrectly concluded that our modified paradigm successfully reduced social engagement in females, if not across sexes. Through the inclusion of four Control groups, however, we have concluded that our findings do not support our hypothesis, and that we cannot endorse this paradigm in its current state. Instead, our negative findings indicate that this protocol requires additional optimization to achieve its overarching goal. Nevertheless, we are sharing our approach and findings so that others with similar procedure goals can learn from our observations, and improve upon our study’s limitations. If we had been able to continue our investigations, we would have next reduced the mild foot shock amplitude, the number of foot shocks, or some combination thereof. Given the group differences in freezing that emerged early during our social conditioning, milder and/or less aversive stimuli could uncover enduring effects of group conditions. To begin though, it was important here that all our groups attained similar starting points to minimize confounds for testing interpretations. We would also advise other researchers to consider optimizing their apparatus and software, if possible, to facilitate consistent analyses of interaction and freezing, if their conditioning/stress component generates freezing. We used one software to administer foot shocks while simultaneously recording and quantifying freezing, and another software to track animal location and identify time spent within specific arena locations, each in separate apparatus. Both software programs excel at their purposes, but each have unique – and unfortunately incompatible – video formats plus requirements regarding lighting, contrast, and background for optimal behavior/location detection.

Another limitation is that we could not evaluate log-transformed corticosterone levels over time without introducing additional confounding stressors to repeatedly sample blood. Daily blood sampling would reveal if our log-transformed corticosterone observations were attributable to specific days, how log-transformed corticosterone levels mapped on to each day’s behavior measure, and/or if log-transformed corticosterone differences are transient or persistent. Our experiments were all performed in sexually naïve mice. We recognize that sexually experienced mice (for example (Newman et al., 2019)) of both sexes might respond, behaviorally and physiologically, in a manner distinct from what we report here. Earlier practices of dividing mice into ‘susceptible’ and ‘resilient’ groups following social stress have more recently fallen out of favor, given categorization of those responses are context-specific, require substantial rodent numbers for optimal data confidence, and neither phenotype can be universally deemed ‘better’ or ‘worse’ (Koolhaas et al., 2017; Lyons et al., 2023). We therefore evaluated all Social Stimulus + Stress mice as a single group. Still, bimodal distribution of social engagement in Social Stimulus + Stress mice looks possible in all but the different sex Social Stimulus + Stress males. Increasing numbers to establish definitive thresholds with sufficient power for phenotypic grouping could prove informative, particularly for the same sex experiment. The same limitations as listed above would still apply, however. Finally, our studies used a brief protocol spanning only four days, intended for high throughput and maximal comparison to other social stress literature. We therefore cannot speak to whether these outcomes persist for weeks or months, timeframes that are ethologically relevant for social learning and memory. Once a short-term protocol has been optimized, duration of effects would be a critical next step in determining validity of the paradigm.

Here, we have reported a novel, albeit unsuccessful, foray into sex-inclusive efforts to develop a controllable, less labor-intensive, and higher throughput social stress paradigm in mice. These experiments have provided utilizable insights regarding how mice of both sexes associate a same-or different-sex target with an aversive experience. We report that these associations seem strongest in males after same sex encounters, whereas females appeared not to form social stimulus associations. Such distinctive responses are probably evolutionarily favorable. While our investigation has generated new information, our modified paradigm did not accomplish what we anticipated. We share our negative findings here to encourage and assist other labs to achieve their own positive outcomes, and to spare them (and mice) the time and effort of duplicating the present work. Development of a sex-inclusive social stress paradigm for mice remains a worthwhile pursuit, as such would be a boon across multiple disciplines.

## Author contributions

JNB, SD, AMJ, LMG designed research. JNB, MMN, IRS, LRS, and LMG performed research. LMG contributed analytic tools. JNB and LMG analyzed data and wrote the manuscript. All authors reviewed and approved the manuscript.

## Acknowledgements

We gratefully acknowledge the mice used in this study, the unrivaled veterinary care by Stan Dannemiller, DVM, MS, DACLAM, and the dedicated work of our vivarium caretakers. Figure 1 was created with BioRender.com (Toronto, ON).

