## Supplemental Material for "Lessons learned while evaluating how pairing aversive experiences with same or different sex social stimulus mice affects subsequent social engagement"

Abbreviated title: Sex effects on social conditioning and behavior (48 characters, 50 character max)

Jasmin N. Beaver^a,b,c^, Marissa M. Nicodemus^a,b^, Isabella R. Spalding^a^, Lauren R. Scrimshaw^a,b^, Sohini Dutta^b,d^, Aaron M. Jasnow^e^, Lee M. Gilman^a,b,c^*

^a^Department of Psychological Sciences, Kent State University, Kent, OH, USA 44242

^b^Brain Health Research Institute, Kent State University, Kent, OH, USA 44242

^c^Healthy Communities Research Institute, Kent State University, Kent, OH, USA 44242

^d^School of Biomedical Sciences, Kent State University, Kent, OH, USA 44242

^e^Department of Pharmacology, Physiology, and Neuroscience, University of South Carolina School of Medicine, Columbia, SC, USA 29209

Author contributions: JNB, SD, AMJ, LMG designed research. JNB, MMN, IRS, LRS, and LMG performed research. LMG contributed analytic tools. JNB and LMG analyzed data and wrote the manuscript. All authors reviewed and approved the manuscript.

Correspondence should be addressed to:

Lee M. Gilman, Ph.D.

600 Hilltop Dr.

209 Kent Hall

Kent State University

Kent, OH, USA 44242

Numbers of:

Figures – 4

Tables – 8

Multimedia – 0

Words for Abstract – 233 (250 word max)

Words for Significance Statement – 102 (120 word max)

Words for Introduction – 750 (750 word max)

Words for Discussion – 2200 (3000 word max)

Acknowledgements: We gratefully acknowledge the mice used in this study, the unrivaled veterinary care by Stan Dannemiller, DVM, MS, DACLAM, and the dedicated work of our vivarium caretakers. Figure 1 was created with BioRender.com (Toronto, ON).

Conflict of Interest: The authors declare no competing financial interests.

Funding sources: This work was supported by R15 MH118705 to AMJ and LMG, and by Kent State University.

**Supplemental Table S1.** Two-way general linear model on cort for mice of both sexes with same sex targets.

| Same Sex Experiment – Log-transformed Serum Corticosterone | | | |
| --- | --- | --- | --- |
|  | **F Statistic** | **p value** | **partial η^2^** |
| Group | F_(4,68)_=7.098 | <0.001 | 0.295 |
| Sex | F_(1,68)_=0.813 | 0.370 | 0.012 |
| Group × Sex | F_(4,68)_=1.207 | 0.316 | 0.066 |

**Supplemental Table S2.** Two-way general linear model on cort for mice of both sexes with different sex targets.

| Different Sex Experiment – Log-transformed Serum Corticosterone | | | |
| --- | --- | --- | --- |
|  | **F Statistic** | **p value** | **partial η^2^** |
| Group | F_(4,70)_=3.841 | 0.007 | 0.180 |
| Sex | F_(1,70)_=8.708 | 0.004 | 0.111 |
| Group × Sex | F_(4,70)_=0.709 | 0.589 | 0.039 |

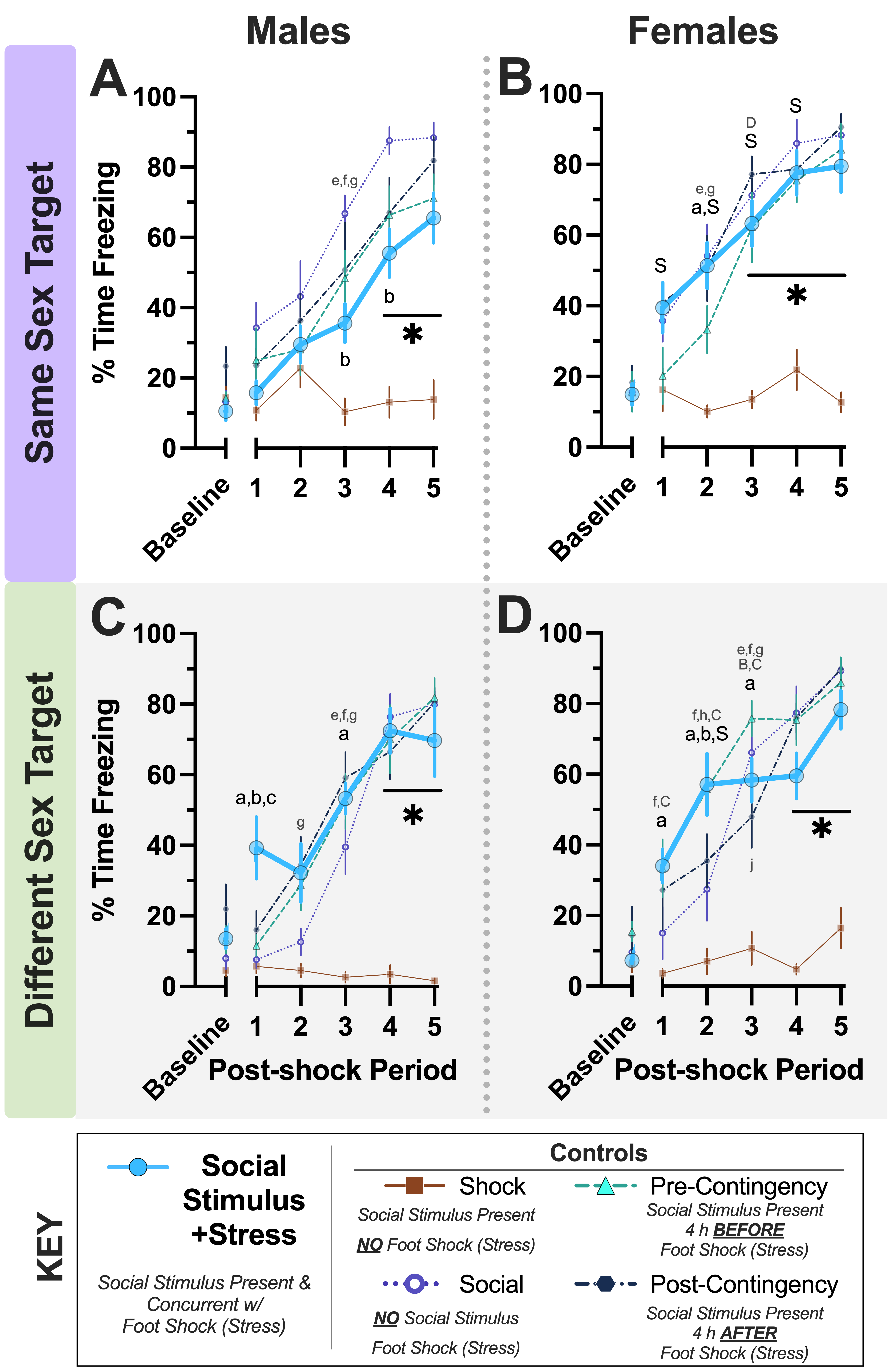

**Supplemental Figure S1.** ***Day 1 acquisition during social conditioning procedure***. Percent time freezing during social conditioning acquisition for mice in same sex (Panels A, B) and different sex (Panels C, D) experiments for male (A, C) and female (B, D) mice. Numbers of mice graphed within Panels A-D in order: Social Stimulus + Stress (n=8, 9, 9, 8); Shock Control (8, 8, 7, 7); Social Control (n=8 for all); Pre-Contingency Control (n=7, 8, 8, 9); Post-Contingency Control (n=7, 8, 8, 8). Average freezing for the first two minutes, prior to commencement of acquisition, is plotted on the x-axis as baseline. The average percent freezing for each 30 second period following each of the five mild foot shocks are thereafter plotted along x-axis (Post-shock Periods 1-5). Alphabetically, within experiment, sex, and time point: ^a^indicates (Shock Control vs. Social Stimulus + Stress) p<0.001, p=0.007, p<0.001, p=0.025, p<0.001, p<0.001. ^b^indicates (Social Control vs. Social Stimulus + Stress) p=0.026, p=0.005, p=0.009, p=0.038. ^c^indicates (Pre-Contingency Control vs. Social Stimulus + Stress) p=0.032. ^e^indicates (Shock Control vs. Social Control) p<0.001, p<0.001, p=0.003, p<0.001. ^f^indicates (Shock Control vs. Pre-Contingency Control) p=0.004, p<0.001, p=0.024, p<0.001, p<0.001. ^g^indicates (Shock Control vs. Post-Contingency Control) p=0.002, p=0.002, p=0.047, p<0.001, p=0.002. ^h^indicates (Social Control vs. Pre-Contingency Control) p=0.043. ^j^indicates (Pre-Contingency Control vs. Post-Contingency Control) p=0.028. ^✱^indicates p<0.001 Social, Pre-Contingency, and Post-Contingency Controls and Social Stimulus + Stress mice vs. Shock Controls. Alphabetically, within experiment, group, and time point: ^B^indicates (Social Controls male vs. female) p=0.005. ^C^indicates (Pre-Contingency Controls male vs. female) p=0.019, p=0.006, p=0.012. ^D^indicates (Post-Contingency Controls male vs. female) p=0.012. ^S^indicates (Social Stimulus + Stress male vs. female) p=0.007, p=0.031, p=0.006, p=0.012, p=0.012. Data graphed as mean ± standard error of the mean.

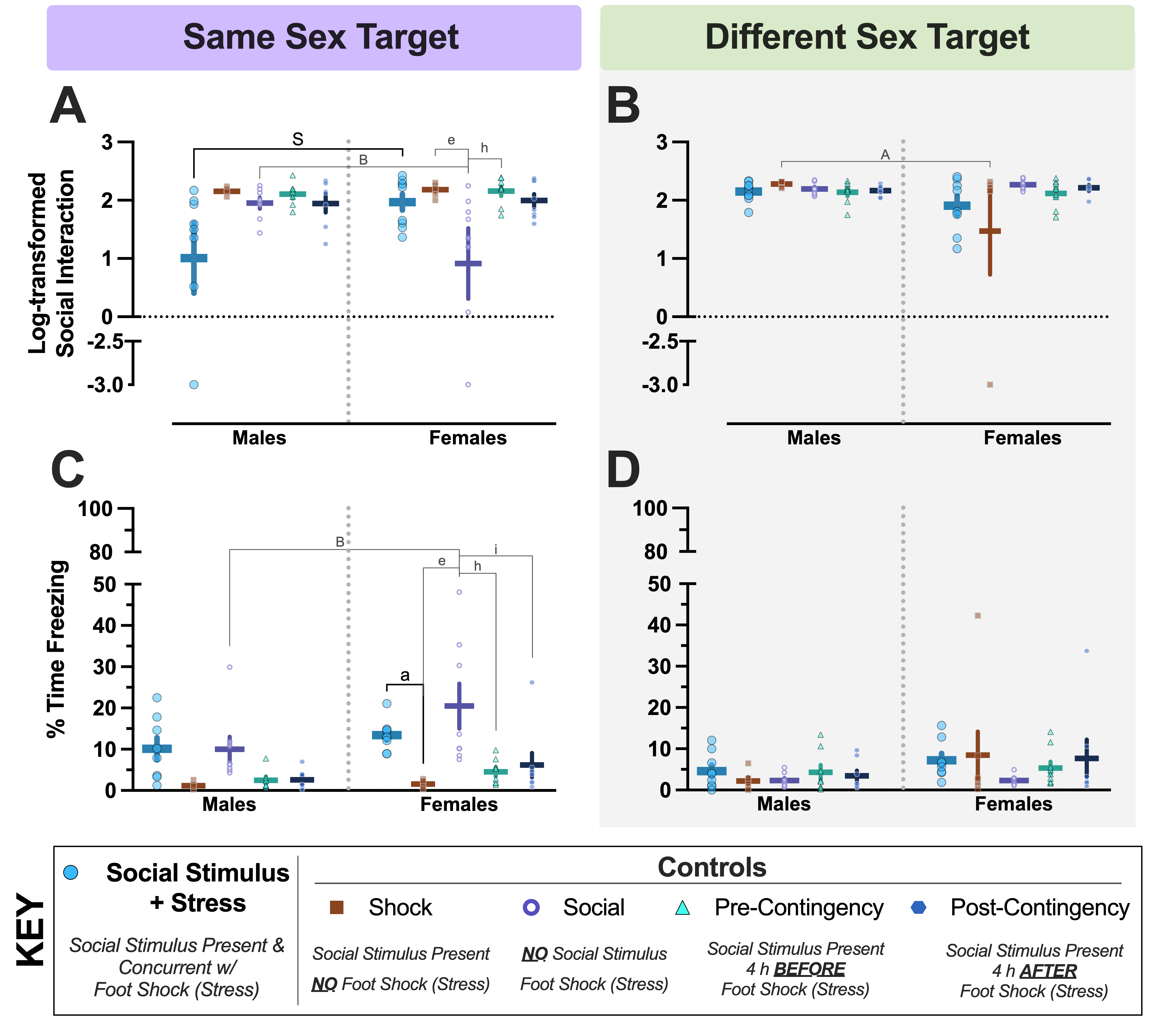

**Supplemental Figure S2.** ***Day 2 social interaction and fear behaviors following social conditioning***. Log-transformed social interaction (Panels A, B) and percent time freezing during post-test social interaction (Panels C, D) data are shown for mice in the same (A, C) and different (B, D) sex experiments. Numbers of mice graphed within each panel, left to right: A) n=8, 7, 8, 7, 7, 9, 8, 8, 8, 8; B) n=9, 6, 8, 8, 8, 8, 7, 7, 9, 7; C) n=8, 7, 8, 7, 7, 9, 8, 8, 8, 8; D) n=9, 6, 8, 8, 8, 8, 7, 7, 9, 7. Y axes for C, D were split to facilitate clearer visualization of the low freezing levels exhibited during social engagement testing. Left to right, top to bottom: ^S^p=0.018, ^B^p=0.013, ^e^p=0.027, ^h^p=0.032, ^A^p=0.023 [authors’ note: we consider this a false positive], ^B^p=0.004, ^a^p=0.008, ^e^p<0.001, ^h^p<0.001, ^i^indicates (Social Control vs. Pre-Contingency Control) p=0.001. Data graphed as mean ± standard error of the mean.

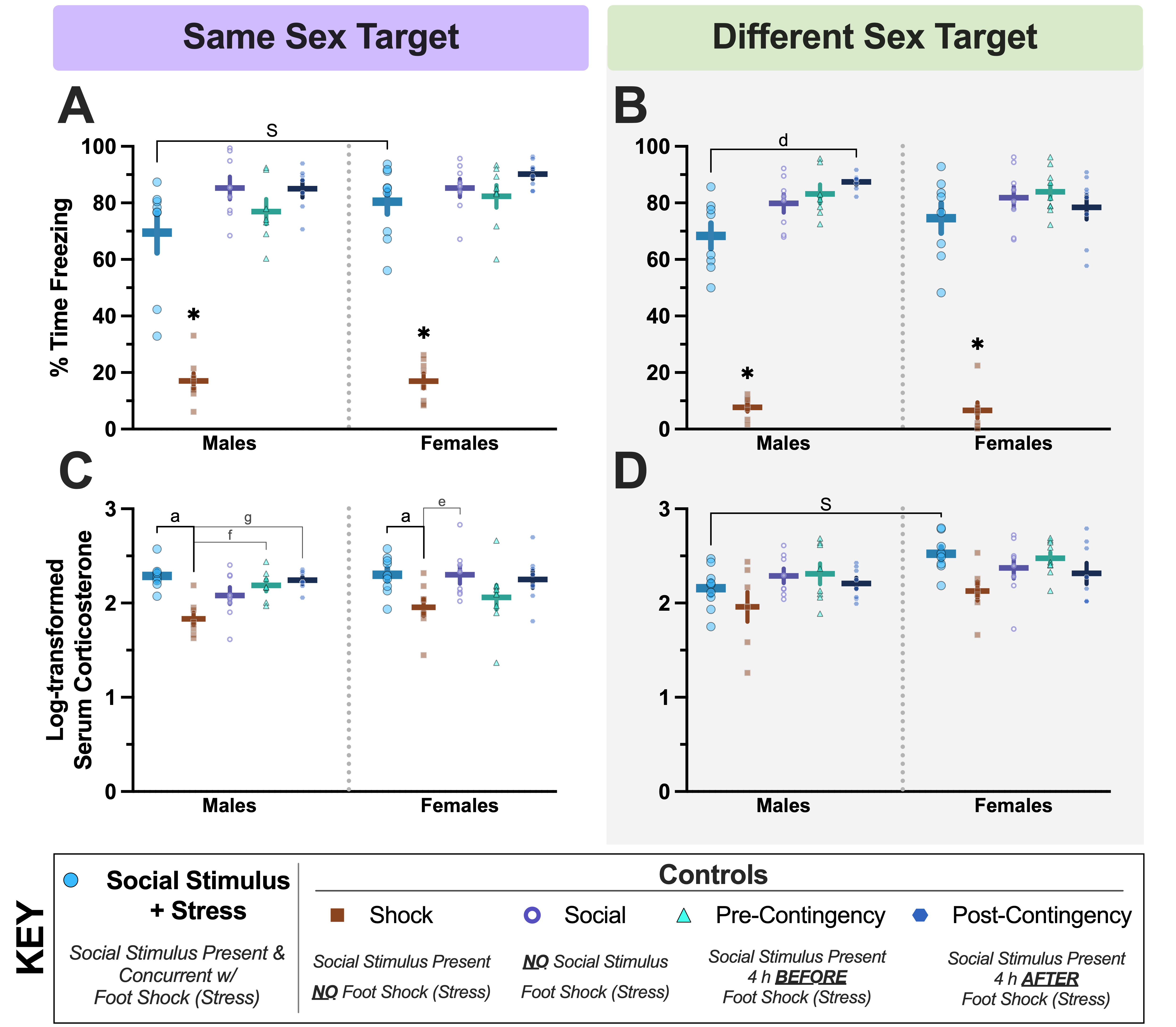

**Supplemental Figure S3.** **Day 3 social conditioning context fear expression averages and subsequent cort levels**. Average context fear behavior (Panels A, B) and cort (Panels C, D) data are shown for mice in all groups in the same sex (A, C) and different sex (B, D) experiments. Numbers of mice graphed within Panels A-D in order, left to right: A) n=8, 8, 8, 7, 7, 9, 8, 8, 8, 8; B) n=8, 7, 8, 7, 7, 8, 7, 8, 9, 8; C) n=7, 8, 8, 7, 7, 9, 8, 8, 8, 8; D) n=9, 7, 8, 8, 8, 8, 7, 8, 9, 8. Left to right, top to bottom: ^S^p=0.049, ^d^indicates (Pre-Contingency Control vs. Social Stimulus + Stress) p=0.004, ^a^p=0.004, ^f^p=0.045, ^g^p=0.012, ^a^p=0.033, ^e^p=0.044, ^S^p=0.007. ^✱^p<0.001 Social, Pre-Contingency, and Post-Contingency Controls and Social Stimulus + Stress mice vs. Shock Control. Data graphed as mean ± standard error of the mean.
